# Liquid plug propagation in computer-controlled microfluidic airway-on-a-chip with semi-circular microchannels

**DOI:** 10.1101/2023.05.24.542177

**Authors:** Hannah L. Viola, Vishwa Vasani, Kendra Washington, Ji-Hoon Lee, Cauviya Selva, Andrea Li, Carlos J. Llorente, Yoshinobu Murayama, James B. Grotberg, Francesco Romanò, Shuichi Takayama

## Abstract

This paper introduces a two-inlet, one-outlet lung-on-a-chip device with semi-circular cross-section microchannels and computer-controlled fluidic switching that enables a broader systematic investigation of liquid plug dynamics in a manner relevant to the distal airways. A leak-proof bonding protocol for micro-milled devices facilitates channel bonding and culture of confluent primary small airway epithelial cells. Production of liquid plugs with computer-controlled inlet channel valving and just one outlet allows more stable long-term plug generation and propagation compared to previous designs. The system also captures both plug speed and length as well as pressure drop concurrently. In one demonstration, the system reproducibly generates surfactant-containing liquid plugs, a challenging process due to lower surface tension that makes the plug formation less stable. The addition of surfactant decreases the pressure required to initiate plug propagation, a potentially significant effect in diseases where surfactant in the airways is absent or dysfunctional. Next, the device recapitulates the effect of increasing fluid viscosity, a challenging analysis due to higher resistance of viscous fluids that makes plug formation and propagation more difficult particularly in airway-relevant length scales. Experimental results show that increased fluid viscosity decreases plug propagation speed for a given air flow rate. These findings are supplemented by computational modeling of viscous plug propagation that demonstrate increased plug propagation time, increased maximum wall shear stress, and greater pressure differentials in more viscous conditions of plug propagation. These results match physiology as mucus viscosity is increased in various obstructive lung diseases where it is known that respiratory mechanics can be compromised due to mucus plugging of the distal airways. Finally, experiments evaluate the effect of channel geometry on primary human small airway epithelial cell injury in this lung-on-a-chip. There is more injury in the middle of the channel relative to the edges highlighting the role of channel shape, a physiologically relevant parameter as airway cross-sectional geometry can also be non-circular. In sum, this paper describes a system that pushes the device limits with regards to the types of liquid plugs that can be stably generated for studies of distal airway fluid mechanical injury.

## Introduction

This paper presents a microfluidic airway-on-a-chip system that expands the range of clinically observed distal lung crackle-associated air-liquid two-phase fluid dynamics that can be recapitulated *in vitro*. The device development work is motivated by the need to recapitulate a broader range of clinical situations including pulmonary edema and mucus obstruction.

The distal lungs are those several generations proximal to the gas-exchanging alveoli, with diameters <2 mm ^1,2^. The epithelium lining these airways is kept moist by a thin annular coating of airway surface liquid (ASL). However, at this length scale the annular coating is susceptible to coalescence into an obstructive liquid plug due to the Rayleigh-Plateau interfacial instability that results from fluids’ tendency to minimize surface tension (Figure 1A)^3,4^. On a large scale, small airway obstruction compromises respiratory mechanics, making breathing more difficult^2,5^. Healthy airways therefore avoid forming obstructive liquid plugs by benefitting from alveolar-produced surfactant proteins that reduce the surface tension at the air-liquid interface between the ASL and inspired air. Diseased airways, however, suffer from compromised ASL surfactant function either due to surfactant dilution or reduced production from dead or dysfunctional cells^6^. This lack of endogenous surfactant favors liquid plug formation, and this is compounded by excessive fluid volume in the lower airways^7^. Liquid plugs then propagate against a pressure gradient, typically applied by inhalation, leaving behind a trailing liquid tail until the plug loses integrity and ruptures (Figure 1A)^8^.

**Figure 1.**
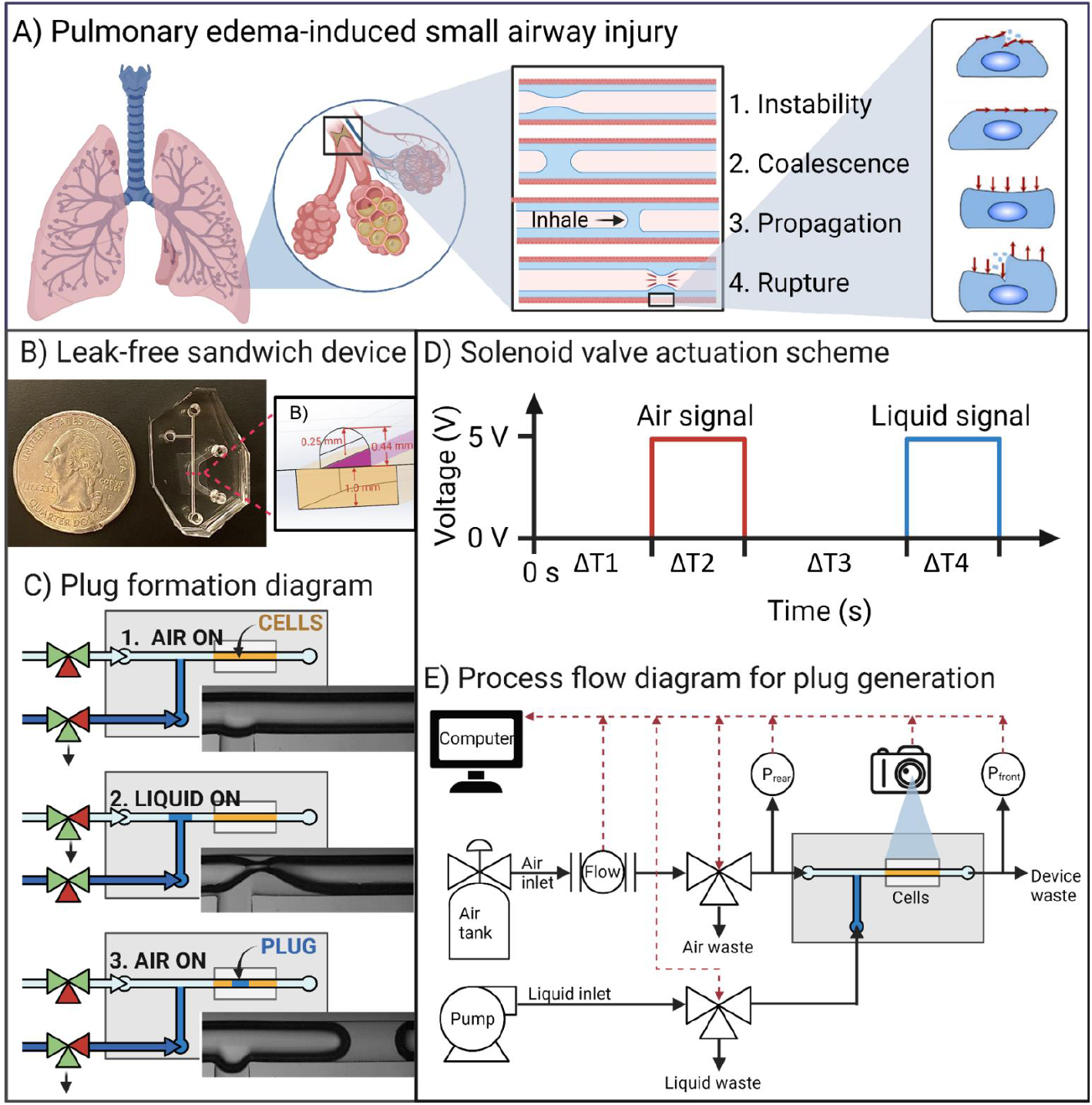
Liquid plug-mediated small airway injury is modeled in a microfluidic device. **A)** Excessive fluid build-up and surfactant dysfunction in the distal airways can lead to fluid-mediated small airway injury. (middle) The air-liquid interface lining the small airways is susceptible to Rayleigh-Plateau instability that causes coalescence into an obstructive liquid plug. Air pressure (for example, during inhalation) pushes the liquid plug until it loses integrity and ruptures, injuring the small airway epithelium. **B)** Microfluidic device image and cross-section showing the design parameters that mimic a distal airway. **C)** Plug formation mimics natural coalescence of liquid plugs by allowing for formation of Rayleigh-Plateau instabilities *in situ* using digitally controlled three-way valves. **D)** Plot of voltage delivered to the air (red) and liquid (blue) valves vs. time (milliseconds). ΔT1, ΔT2, ΔT3 and ΔT4 are arbitrary, tuned electronically through LabView. **E)** Process flow diagram for plug generation scheme.

Additionally, mucus and airway surface liquid in patients with obstructive lung diseases, such as chronic obstructive pulmonary disease, can be far greater than in healthy individuals^9^. In these patients, mucus in the small airways can form obstructions called mucus plugs that may compromise airway reopening in collapsed small airways^10,11^. Obstructive lung diseases, especially chronic obstructive pulmonary disease (COPD) and cystic fibrosis (CF), are associated with intrinsic airway narrowing, increased airway smooth muscle tone, and potentially increased airway stiffness that may preclude true collapse due to the lack of elasticity of the airway walls^12,13^. Importantly, however, airway elasticity is not required to initiate a liquid obstruction due to Rayleigh-Plateau instability, as demonstrated by our recent computational modeling of airway closure in a rigid cylinder model^7^. Therefore, individuals with rigid airways due to muco-obstructive lung disease may experience obstruction without collapse, necessitating studies of viscous effects in the context of rigid airways.

However, for both surface tension and viscosity effects, the cellular and molecular mechanisms underpinning fluid mechanical stress-mediated cellular injury are poorly understood due to a lack of adequate model systems. The prevailing approach has been computational due to the tunability of fluid dynamical modeling systems^14^. These have significantly advanced the field’s understanding of the fluid dynamics and stresses imparted onto the epithelium during liquid plug formation, propagation, and rupture, and allow tuning of experimental variables such as fluid viscoelasticity and airway flexibility^7,11,14–17^. However, computational models cannot predict cellular responses to these well-studied stressors, which is critical to understanding how these forces affect pulmonary pathophysiology. Therefore, an experimental model system incorporating cells is required. Animal models allow complex airway physiology that is often advantageous, but here they offer severely limited control of airway and fluid properties, such as plug viscosity, surfactant content, size, speed, and frequency in individual airways. An *in vitro* approach would enable such precise control, but the complex fluid dynamics of this system preclude the use of commercially available *in vitro* culture systems, such as Transwells^18^ or commercial microfluidic devices^19^.

Therefore, our group previously fabricated a two - inlet, two-outlet microfluidic lung-on-a-chip to model liquid plug propagation and rupture above a monolayer of airway epithelial cells^8^. This model enabled us to probe the effects of surfactant concentration and channel hydrophobicity on cell viability^20^ and plug properties^21^ (i.e., length and speed), respectively. We showed that surfactant decreases the pressure force required to propagate a liquid plug and thus reduces wall shear stress that would injure cells^22^. Additionally, we found through joint experimental and computational studies that surfactant reduces the formation of Rayleigh-Plateau instabilities and therefore reduces the incidence of plug formation^22^. We termed these injurious liquid forces that impart lethal shear stress to airway epithelial cells on the walls “fluid mechanical stressors”^8,20,21,23,24^. Our results suggest that fluid mechanical stress could play a significant role in cellular-level pulmonary pathophysiology, especially in edematous and muco-obstructive lung diseases ^4^. However, our design suffered from poor control over plug generation. The time interval between plugs was difficult to control^21^ and the device could not reproduce identical plug size and length for long time periods (i.e., hours to days rather than minutes, unpublished observation). The two-outlet design made manipulation of viscous liquids particularly difficult due to constant changes in the relative resistances of the two different outlet paths. These technical limitations reduced the stability and range of liquid plug conditions that could be realized. Indeed, controlling plug properties is critically important to experimental robustness and reproducibility, and to the investigation of viscous and surfactant laden fluids.

Therefore, we present and characterize a novel liquid plug generating microfluidic lung-on-a-chip system with semi-circular cross-section microchannels coupled with peripheral valves and computerized controls to enable a broader range of small airway liquid plug formation, propagation, and rupture in a reproducible, well-controlled process with live monitoring of pressure, air and liquid flow rates, and plug length and speed *via* high-speed microscope-mounted camera. Programmable three-way valves enable arbitrary, automated control of plug length, speed, and interval between plugs for unprecedented control of experimental conditions. Then, we characterize capillary resistance that is exacerbated by high surface tension, here caused by low surfactant concentration. In our integrated system we show that this effect may be significant physiologically important, highlighting its necessity in future fluid dynamics studies. Additionally, we show that viscosity effects may exacerbate liquid plug induced forces and their pathophysiologic significance. Finally, the device incorporates confluent primary airway epithelial cells with novel insight on the role of the semi-circular channel geometry. Together, these technological improvements represent a significant advance towards understanding fluid mediated distal airway injury with recapitulation of pathophysiologic surface tension and viscosity effects.

## Materials and Methods

### Microfluidic device fabrication

Microfluidic device fabrication followed the process depicted in Figure 2A. In brief, 3D models for the top and bottom microfluidic channels were custom designed in SolidWorks. Negative molds were micromilled into acrylic using paths designed in Fusion360 (AutoDesk) with a 1 mm square end mill and 200 μm and 100 μm ball nose end mills (Performance Micro Tool and Harvey Tool). Positive molds were cast into the acrylic negative mold with polydimethylsiloxane (PDMS) in a 5:1 weight:weight ratio with SU-84 crosslinking agent. The PDMS-crosslinker was poured onto the acrylic mold, placed under vacuum for 20 minutes to remove air bubbles, and baked at 65 °C overnight. The PDMS was then separated from the acrylic mold and further baked at 120 °C overnight. Finally, the PDMS positive mold was used to cast devices in PDMS. PDMS was mixed 1:10 wt:wt with SU-84 crosslinker and poured over the PDMS positive mold. This was placed under vacuum for 20 minutes, and then baked for 2 hours at 65 °C. The PDMS devices were then separable from the PDMS mold. Devices were cut out, and holes were biopsy punched (1.5 mm diameter). Membranes were fabricated from Transwell inserts (Corning 353102) with pore size 1 μm. Membranes were cut out of the plastic Transwell insert and treated with 5% volume/volume (3-Aminopropyl)-triethoxysilane (APTES) solution in distilled water at 80 °C for 20 minutes. Temperature during the surface treatment was maintained by periodic verification with a laser thermometer. Following the surface treatment, membranes were hung to dry in the fume hood overnight. Finally, devices were assembled using plasma oxidation. Membranes and top channel were treated with oxygen plasma for 30 seconds at 120 volts and the membrane was bonded to the top channel. Then, the top channel and membrane together were treated with the bottom channel (30 seconds, 120 V) and the top and bottom channels were assembled. The finished device was post-baked at 65 °C for an additional 2 hours to solidify the bond.

**Figure 2.**
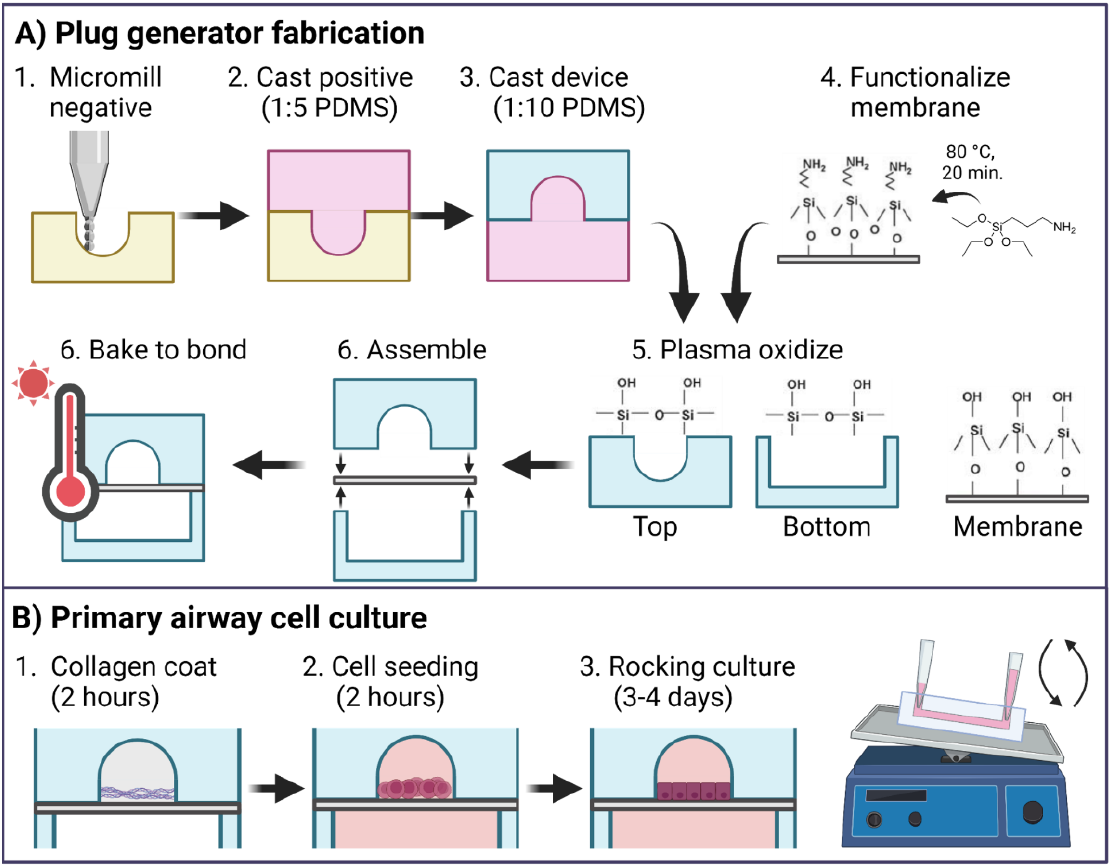
Device fabrication protocol. **A)** Microfluidic liquid plug generating device fabrication strategy. **B)** Primary small airway epithelial cell culture strategy in the liquid plug generator. Medium perfusion is accomplished by gravity-driven flow on a plate rocker, avoiding the requirement for pumps or tubing.

### Cell culture

Cell culture was executed as summarized in Figure 2B. Assembled devices were sterilized under UV light for 20 minutes prior to use. Type I rat tail collagen (Corning 354236) was suspended at 26 μL/mL in cold 1X PBS (without calcium or magnesium) that was previously adjusted to pH 9 with sodium hydroxide to avoid collagen gelation and sterilized by membrane filtration with 0.22 μm pore size. The top channel of the device was filled with the collagen suspension and was allowed to incubate for 90 minutes in the cell culture incubator (37 °C, 95% humidity, 5% CO_2_). Following incubation, the collagen solution was replaced with cell culture media and the liquid entry point was plugged to prepare for cell seeding. The entrance and exit to both top and bottom channel were given cut-off P1000 pipette tips to serve as media reservoirs and to deliver cells and media. Primary small airway epithelial cells were purchased from Lonza (CC-2547) and expanded in Lonza’s small airway growth medium with BulletKit supplements as instructed (CC-3118). For device seeding, cells were passaged at 60-80% confluence according to Lonza instructions using Lonza Reagent Pack (CC-5034). Small airway cells were then seeded into devices at 3-4 million cells/mL, 30 μL/channel, by adding the cells to the top channel pipette tip to avoid lethal excess shearing. Cells were allowed to flow across the top channel and settle evenly. The devices were incubated for 2 hours to allow cell attachment, and then 100 μL of cell culture media was perfused across the top channel to remove unattached cells. Devices were cultured under liquid-liquid conditions for 2 days following seeding with daily media exchanges. Devices were placed on a wave plate inside a cell culture incubator (37 °C, 95% humidity, 5% CO_2_). The wave plate rocked 1°/second until the maximum angle of the wave plate, then rocked back. Medium was changed every 24 hours.

### Plug generation

Valve control was achieved by programming LabVIEW (National Instruments) to send arbitrary signals to an Arduino microcontroller that communicated with two 3-way solenoid valves (LFN series, Lee Company) that had a 20 ms average response time. The air flow was delivered from a compressed air tank and first passed through a mass flow meter (Bronkhorst for Elveflow) that was density, temperature, and pressure-independent due to its basis in the Coriolis principle. Air then entered the 3-way valve, split to a pressure meter (Omega Engineering, PX409-001DWUUSBH), and finally entered either to the device or to the vent. The computerized implementation also allowed simultaneous data acquisition for pressure data. The liquid was fed from a flow-rate controlled syringe pump (NemeSys) and passed through the 3-way valve and pressure meter into either to a drain line or into the device. The device outlet is attached to a 2-inch drain tube at atmospheric pressure that ends in a conical vial to collect waste. The liquid plug generator was mounted on a brightfield microscope (Olympus) to capture images of plug propagation and rupture. The images were captured by a Phantom high speed microscope camera (Phantom VEO-E 310L by Vision Research). Plug generation and propagation was captured at 500 frames/second at a resolution of 832×400 pixels. Plug speed was defined using the 2-point instant active measurement. The first point being the end of the plug at rest and the second that same edge of the plug before it left the camera frame.

### Cell counting and statistics

Following plug exposure, devices were incubated for 10 minutes with a mixture of live cell-permeable NucBlue (Invitrogen R37605) and live-cell impermeable Sytox Green (Invitrogen S7020) followed by rinsing with Hank’s Balanced Salt Solution. Membranes were cut out from devices and mounted on glass slides. Images were captured, using identical settings across conditions, on a Leica DMI8 Microscope. Image-based quantitation of cells from resultant immunofluorescent stains was performed using the ImageJ segmentation plugin CellPose ^25^.

### Simulation of plug propagation

The simulation of the multiphase flow involving gas and liquid phases is carried out using a single-field formulation, which deals with both phases simultaneously. The liquid plug rupture is assumed to be an axisymmetric phenomenon. The simulations are conducted using an open-source code called *basilisk*^26^, which implements a volume-of-fluid (VOF) method. This method utilizes a second-order finite volume discretization in space and a second-order pressure-correction projection method for time integration. A semi-implicit approach is utilized, which discretizes the viscous terms implicitly and the convective terms explicitly using the Bell–Collela–Glaz advection scheme^27^. The VOF method belongs to the front-capturing methodologies and employs a fraction field to define the variable material properties in the single-field approach. The fraction field denotes the volume fraction of liquid in a computational cell and ranges between 0 and 1 across the liquid-gas interface.

To solve the equation, a piecewise-linear geometrical VOF method is employed, which represents the interface as a piecewise linear function within each computational cell. A staggered approach is utilized to achieve second-order accuracy in time. In the staggered grid, the pressure and velocity fields are located at the cell centers and faces, respectively. Scalar fields such as volume fraction and material properties of the gas and the liquid are also located at the cell centers. Once the local unit vector at the interface is computed using the mixed Youngs-centered method, material properties are set in each phase. The surface-tension term, which appears in the momentum equation of a single-field model, is discretized. The approach by Popinet is used^28^, which combines a height-function estimator for interface curvature and a balanced-force surface-tension discretization. This combination reduces numerical parasitic currents significantly and yields accurate validation for challenging benchmark cases in pulmonary flows^7,29^.

## Results

### Device design and fabrication strategy

To replicate the distal airways to capture physiologically relevant fluid dynamics, we modeled the terminal bronchioles that are the most distal airways with bulk air flow. Their diameter ranges from 0.5-0.1 mm ^30^. We used a micro mill to design a semi-rounded channel that would approximate the shear stress profile of the airways in a semi-circular conduit^31^. This design allowed us to culture airway epithelial cells on a porous substrate while modeling the rest of the airway channel with rounded geometry. To determine the air speeds required in our device, we utilized general calculations for air speeds based on airway generation developed by Weibel^30^. The approximate velocity of airflow at airway diameter 0.05 cm in a healthy adult is calculated as follows:

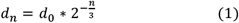

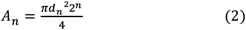

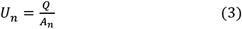

*d*_*n*_ = airway diameter at generation n [cm], *d*_0_ = tracheal diameter [cm], *n* = airway generation, *A*_*n*_ = cross-sectional area of generation n airway, *Q* = flow rate of air into the lungs [mL/s], *U*_*n*_ = air velocity at generation n airway [cm/s]

The resultant Reynolds number (*Re*) can be calculated as follows:

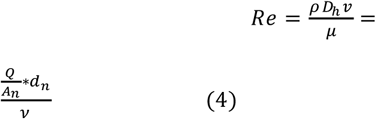

where ρ = density [kg/m^3^], D = hydraulic diameter [m], μ = dynamic viscosity [kg/m-s], v = kinematic viscosity [m^2^/s]

Assuming a round airway of diameter 0.5 mm, a breathing rate of 500 mL/s, tracheal diameter of 2 cm, and kinematic viscosity of air 0.15 m^2^/s, the target airway generation is 16 and the velocity is 3.95 cm/s and *Re* = 1.31. Therefore, airflow is laminar in this region. We aimed to replicate this air speed in the channel while incorporating physiologically rounded airway geometry. Therefore, we designed a channel with hydraulic diameter of 0.46 mm and cross-sectional area 0.193 mm^2^ (Figure 1B)). This semicircular geometry closely approximates a round tube with diameter 0.5 mm and cross-sectional area 0.196 mm^2^ (Table 1). Further, the measured air velocity in the microfluidic channel confirms that air velocity approaches the Weibel calculation (Table 1).

### Consistent, long-term plug generation

While previous plug generating microfluidic airway devices have been reported, this device enables stable plug generation over more extended time scales (minutes to hours) and with an expanded range of liquid properties. To pilot the capabilities of our plug generation scheme, we tested a simplified version of the device consisting only of the top layer bonded to a glass slide using plasma oxidation. Under these conditions, however, the channel may be more hydrophobic than the full device where cell culture media and cells have been allowed to incubate in the channel and adhere proteins for several days prior to plug generation. Therefore, to match the channel properties to our final cell culture device, we coated the simplified glass slide-bound channel with 5% wt/vol bovine serum albumin for 10 minutes to allow protein adsorption. While it is difficult to measure the hydrophilicity of our channel due to its geometry, BSA coating is known to promote hydrophilicity of PDMS surfaces after plasma oxidation^32^, and it is a major component of the cell culture medium for small airway cells that incubates with devices during culture. We observed that 5% BSA treatment restored plug edges to a concave curve as opposed to convex, confirming the increased hydrophilicity^21^ (unpublished observation). High-speed movies of the moving liquid plug, however, still show significant contact angle hysteresis especially for the advancing meniscus. While cell surfaces are often assumed to be highly hydrophilic, they can have relatively high contact angle and surface heterogeneity particularly for air-exposed tissues such as the amnion^33^, various gastrointestinal mucous-coated surfaces^34–36^ and the air-exposed corneal epithelium^37^. Although measurements of pulmonary mucosa contact angle in the small airways are limited by the tissue’s geometry, several tracheal studies have suggested that under healthy conditions the lung mucosa is relatively hydrophilic. However, the contact angle is influenced by surfactant and mucous content, which are severely compromised in the setting of various pulmonary diseases such as acute respiratory distress syndrome and cystic fibrosis^38^. Indeed, multiple studies have demonstrated that contact angle hysteresis is observed on pulmonary epithelia in the presence of surfactant, suggesting that while surfactant improves wetting it does not necessarily enable complete wetting that eliminates hysteresis, as is the case in the corneal epithelium that is also air-exposed^37,39, 40^. Therefore, our observations of contact angles and contact angle hysteresis have physiological relevance in a disease context.

We designed a plug generation process that automates plug generation and enables consistent generation of controlled-volume plugs (Figure 1C, E). The device consists of a straight air channel perpendicular to a liquid injection channel. Conveniently, air and liquid sources (gas tank and syringe pump, respectively) were allowed to generate a constant flow rate. Air and liquid inlets were controlled by three-way electronically actuated valves that diverted air and liquid from the microchannel inlets upon applying a voltage. During liquid injection, air flow was stopped to prevent premature movement of the liquid bolus, another capability not possible with our prior device. After liquid loading was complete, liquid inlet was closed and the air flow returned to the channel, enabling propagation of the liquid plug. Control of valve actuation and therefore air and liquid inlets was achieved using LabView fed to an Arduino microcontroller. Controllable parameters are air flow rate, liquid flow rate, and T1-T4. Liquid plugs were generated according to the strategy described above, for PBS with a liquid flow rate of 600 μL/min and air flow rate of 42 μL/min. ΔT1 = ΔT2 = ΔT3 = 500 ms; Cycle time = 5 s.

This design enabled highly consistent plug generation across over 300 plugs over a 25-minute timespan. Notably, longer timespans are possible in our system, but not necessary for demonstration of our application. We quantified plug properties by measuring the pressure differential across the device over time (Figure 3A). The pressure difference drops back to zero when the plug ruptures. The initial spike in the pressure curve (Figure 3A, circle) represents the extra force required to overcome transient resistance to initiate movement of a stationary plug in a relatively low-wetting or high surface tension context, discussed further in the following sections^41^. The subsequent flat region coincides with plug propagation and represents the energy necessary to move the plug against dynamic viscous and capillary resistances. We found that the device reproducibly generated over 300 plugs with a mean initial spike pressure of 0.235 +/- 0.048 psi (Supplemental Figure 1). Furthermore, the time interval between plugs was normally distributed around a mean of 5.0 seconds (the set point) with only 3/300 plugs falling outside the range of 4.5-5.5 s and 256/300 falling virtually exactly at 5 seconds. Taken together, these metrics demonstrate the reproducibility and long-term stability of the model system for generating liquid plugs in a physiologically relevant set of conditions. These features are critical to future studies that require longer term plug application. Additionally these quality control parameters have not been reported in previous plug models in lung-on-a-chip.

**Figure 3.**
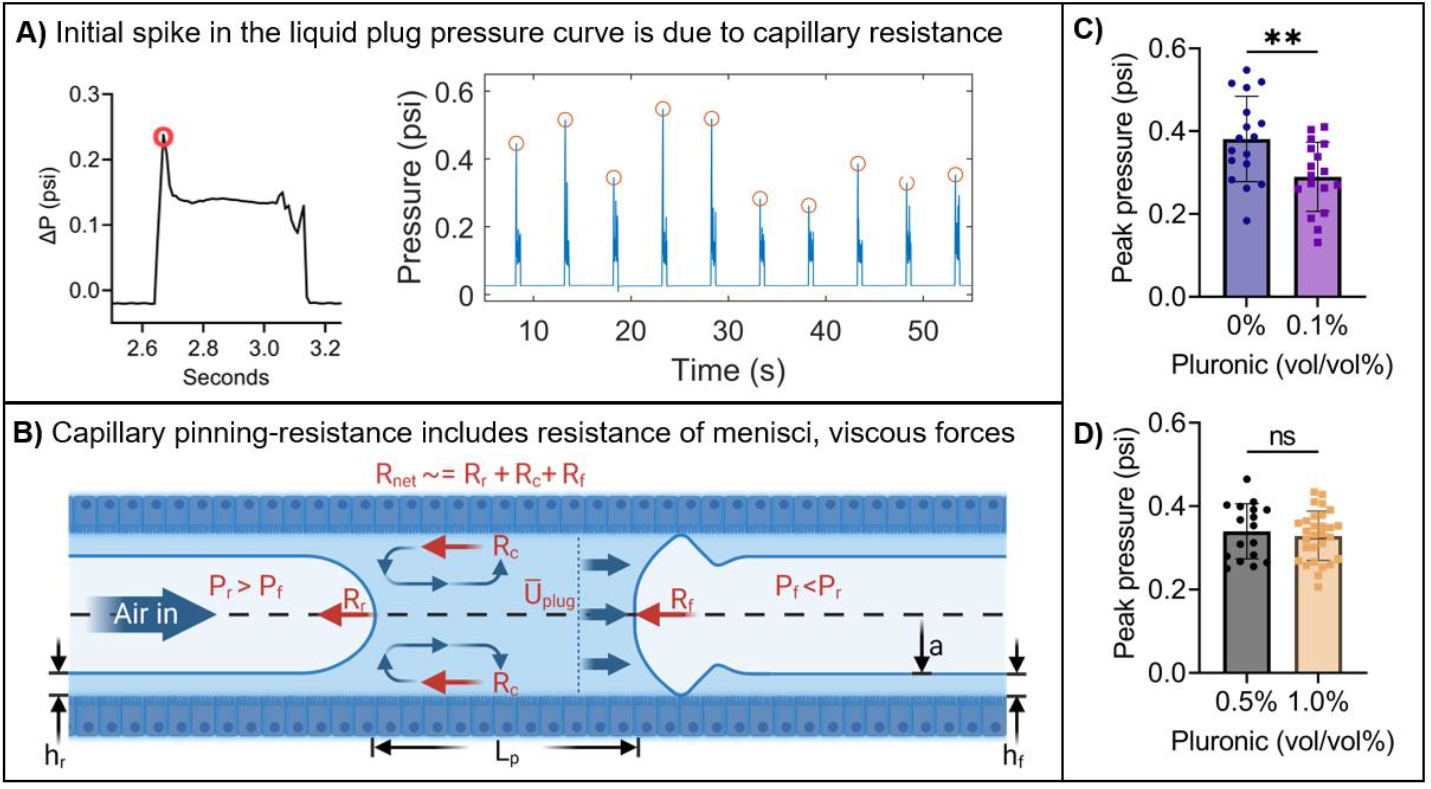
**A)** Pressure vs. time plot from MATLAB, red circles indicate the peak pressure for each plug identified by our custom MATLAB script (supplemental data) **B)** A force balance can approximate the resistance to motion of a liquid plug in the Stokes flow regime. Upon initiation of a driving pressure gradient (P_r_ > P_f_), the initial motion of the liquid plug must overcome viscous (R_c_) and capillary resistance (R_r_, R_f_). Viscous resistance arises from shear stress at the liquid-wall interface while capillary resistance arises from the requirement for movement of the concave air-liquid meniscus at both the front and rear of the liquid plug. Capillary resistance depends on surface tension meaning that significant alterations to surface tension reduces net resistance to both plug movement initiation and steady-state propagation **(C)**. However, once sufficient surfactant is present, further reduction is not achieved due to diminishing returns **(D)**. **p<0.001, student’s t-test

### Surfactant reduces static capillary pinning

Next, we utilized the enhanced capabilities of our model to demonstrate the effect of surfactant on the capillary pinning of a liquid plug at the initiation of plug motion. Static plugs form when air flow halts in the small airways while the lungs transition from exhalation to inhalation. Then, inhalation initiates movement of static plugs. Compared to plug propagation and rupture, initiation of plug motion in the distal airways has not been studied in detail despite its potential to create substantial stress due to capillary pinning in low wetting conditions. Capillary pinning refers to the phenomenon produced by resistance under low-wetting conditions that is typically caused by high surface tension between the two fluids pinned on a solid substrate along a triple-phase contact line (see pressure spike, Fig. 3A). In the lungs, the air-liquid meniscus of a liquid plug has high surface tension due to surfactant dysfunction that results in pathologically thin airway surface liquid coating ^42–44^. This high surface tension, and resultant poor wetting, increases resistance when the plug is static, resulting in capillary pinning. Other sources of resistances are due to the dynamics of the plug. Dynamic resistance is comprised of three major components: the front and rear air-liquid menisci and the interface between the liquid and solid (Figure 3B). The menisci impart the bulk of resistance forces when the plug is static (static equilibrium is not shown in Figure 3B). Upon reaching the critical pressure force needed to overcome this pinning, the force needed to move the plug rapidly declines to a steady value owing to lower dynamic resistance opposed by capillary forces including both menisci and the liquid-solid shear stresses^41,45,46^.

Figure 3B describes these forces. The figure depicts an axisymmetric liquid plug propagating from left to right (see blue arrows labelled with average propagation speed Ū_plug_) in a circular pipe. The pipe wall in the sketch is assumed to be entirely wetted. An air finger at the read plug front is pushing the liquid plug with a pressure P_r_ (see blue arrow labelled “Air in”) and the quasi-spherical interface at the tip of the air finger is associated to a capillary resistance that relates on the curvature of the meniscus and the surface tension between liquid and gas. Such a rear-meniscus resistance is denoted as R_r_ in the sketch. At its leading front, the liquid plug is pushing a second air finger whose core pressure is P_f_, lower than P_r_. This second air finger is compressed, which leads its interface to form a capillary wave near the pipe wall. Consequently, the curvature of the front meniscus for the liquid plug is significantly different from the one of the rear meniscus. Hence, the capillary resistance of the liquid-plug front meniscus R_f_ is different from R_r_. Such two capillary resistances depend on the surface tension, on the propagation velocities of the liquid-gas fronts, as well as on the viscosity of the liquid. In fact, the corresponding physics is governed by their capillary number, which is the ratio between the meniscus propagating velocity, times the liquid viscosity over the surface tension. Finally, for a fully-wetted pipe, a third resistance adds up in order to take into account the viscous resistance at the pipe wall. This core-flow resistance R_c_ depends on the liquid viscosity, the pipe radius, and the plug length L_p_, i.e. the minimum distance from the rear to the front meniscus (distance on the axis for an axisymmetric liquid plug). More details about the specific expressions of such resistances in a Stokes flow approximation is reported in the supplementary material and explained in details in Bahrani et al. (2022)^14^.

Our previous work showed that capillary pinning results in transiently increased capillary resistance that manifests as a spike in pressure differential at the initiation of static plug motion, in a shape mimicking our pressure vs. time curve^14^, Supplemental Equations 1-3. Therefore, pinning effects that create transient excess resistance to flow may be physiologically relevant.

However, previous microfluidic lung-on-a-chip devices have not enabled detailed investigations of the role of transient static-to-dynamic forces in the motion of plugs. Here, we demonstrate this capability by showing that surfactant-free plugs experience measurable pinning force that is quantitatively reduced by the addition of surfactant. We quantified capillary pinning forces by measuring the pressure spike that appears at the initiation of plug motion in our device (Figure 3A, circled). To demonstrate the utility of this quantitative property of plug pinning, we compared the pressure spike magnitude induced by plug generation with pure distilled water compared to distilled water containing 0.1 vol/vol% Pluronic, a polymer-based synthetic surfactant. Plugs were generated at identical air and liquid flow rates of 48 μL/min and 100 μL/min respectively. We also compared the effect of 0.5% vs. 1.0% Pluronic under the same air flow rate but 200 uL/min because the high surfactant concentration made plug formation more difficult. The liquid channel was opened for 500 milliseconds (ms), then 750 ms with both channels closed, then air channel opened for 250 ms. Then, both valves were closed for the remainder of the total 5 second plug generation cycle. We quantified the pressure difference across the channel during the initial pressure spike for both conditions using a custom MATLAB script that identified the maximum value in the pressure curve for each plug (Figure 3A, circles). We also obtained images from high speed camera images showing contact angle hysteresis at the advancing but not trailing menisci.

We found that the initial pressure spike immediately before the plug begins moving was significantly reduced with the addition of surfactant compared to no surfactant, but not when surfactant was already present (Figure 3C, D). Specifically, the peak spike was reduced from 0 to 0.1% pluronics, but not from 0.5 to 1.0% pluronics. This finding suggests that surfactant’s improvements have diminishing benefit to the reduction in pinning forces, an observation we have previously found computationally^22^. We found that less pressure was required to overcome pinning and move the liquid bolus compared to the no-surfactant condition. This result is consistent with other studies showing that surfactant reduces pinning and capillary resistance^47,48^. Additionally, our previous experimental and computational studies that showed surfactant reduces the wall shear stress of moving liquid plugs, a reflection of reduced resistance ^21^. We also note that while cell surfaces are generally hydrophilic, surface heterogeneity that can form during disease leads to contact angle hysteresis and contact line pinning^49,50^. Our device introduced a novel capability to study static pinning forces, a potentially significant pathophysiologic effect, in a lung microenvironment *in vitro* with unprecedented control.

### Viscosity impacts plug speed, pressure drop *in vitro* and *in silico*

Small airway surface liquid can become hyperviscous due to excessive mucins, dead cell debris, pathogen colonization, poor ion balance, and other factors^51^. Various clinical and experimental evidence suggests that small airway hyperviscosity contributes to airway collapse and compromises overall lung compliance^9,11,16,52^. Therefore, we modeled the effect of increased liquid viscosity on plug propagation speed. We chose viscosity values based on measurements of airway surface liquid collected from primary COPD patient airway epithelial cells with varied severity of chronic obstructive pulmonary disease, which is expected to reasonably replicate patient-derived sputum properties^9,53,54^. We generated liquids of increasing viscosity by dissolving PBS with increasing concentrations of 500k Dextran. Experimental viscosities were measured on a rheometer. The control PBS had an average viscosity of 1.0 cP, the healthy individual mimetic, 6.1 cP, healthy smoker mimetic, 12.6 cP and COPD smoker mimetic fluid 628.5 cP^9^.

Varied-viscosity plugs were propagated through microfluidic devices using the same plug generation scheme for each viscosity. First, both channels were closed for 3.5 seconds (from start time until T1, Figure 1D). Then liquid would enter the channel for 0.5 seconds (ΔT1), both channels were closed for 0.5 seconds (ΔT2) then followed by air for 0.5 seconds (ΔT3). Air flow rate was set to 50 +/- 2 μL /min and liquid flow rate was set 100 μL/min. With these conditions we found that fluid viscosity was inversely related with the speed of plug propagation (Figure 4A) as predicted by previous experimental (cell-free) and computational studies on the effect of plug speed and viscosity^11,14^. Next, we simulated plug propagation with varying viscosities that approximated our experimental conditions. Simulated plug propagation speed trended downward with increased viscosity in line with our experimental findings (Figure 4B). The propagation velocity of the liquid plug U_plug_, computed as the mean of rear and front meniscus velocities, is evolving with time. As the resistances R_r_, R_f_, and R_c_ all depend on the liquid viscosity, the propagation velocity of the liquid plug will be strongly affected by μ. Moreover, as R_r_ and R_f_ involve power laws of the liquid viscosity via their corresponding capillary number, the propagation velocity of the plug is supposed to carry a power-law trend. This is confirmed by Figure 4B, where the least-square best fit of the maximum velocity of the liquid plug suggests that max(U_plug_) ∼ μ^-2.7^. Similarly, the average propagation velocity computed in our numerical simulations scales like <U_plug_> ∼ μ^-3^. Additionally, absolute plug speed matched between experiment and computation within an order of magnitude (Figure 4 A, B). It is important to note that the pressure vs. time curve did not include the initial pressure spike because simulations assumed a uniformly distributed wetting film at the wall that would negate wetting-related resistance. However, the pressure curve shape was qualitatively similar to our data (Figure 4C). Additionally, the pressure curve demonstrates that more viscous plugs experience delayed rupture (Figure 4C). The dashed lines in Figure 4C denote the time evolution of the liquid plug length Lp (minimum distance between rear and front menisci) for five liquid plug viscosities μ ranging from 2.16 cP to 12.6 cP. All liquid plugs are initialized with the same length L_p_(t=0) and their rupture time t_r_ at which L_p_(t_r_) = 0 increases with the fluid viscosity. Upon plug propagation, the axisymmetric simulations predict a wall pressure excursion Δp=max(p(r=a))-min(p (r=a)) (solid lines) that is in good agreement with our experimental measurements. Viscous plugs also have greater maximum wall shear stresses when they do rupture, implying that increased viscosity may have a more injurious impact on airway epithelial cells (Figure 4D). Similarly to Figure 4C, Figure 4D correlates the liquid plug dynamics with the maximum shear stress at the wall τ_max_. The typical post-rupture peak of τ_max_ is captured by the numerical simulations, which peak significantly increases with μ up to 4.31cP and tends to saturate beyond such a liquid viscosity. These *in silico* findings corroborate other computational studies of viscosity in liquid plug formation, propagation and rupture^11,14, 29^.

**Figure 4.**
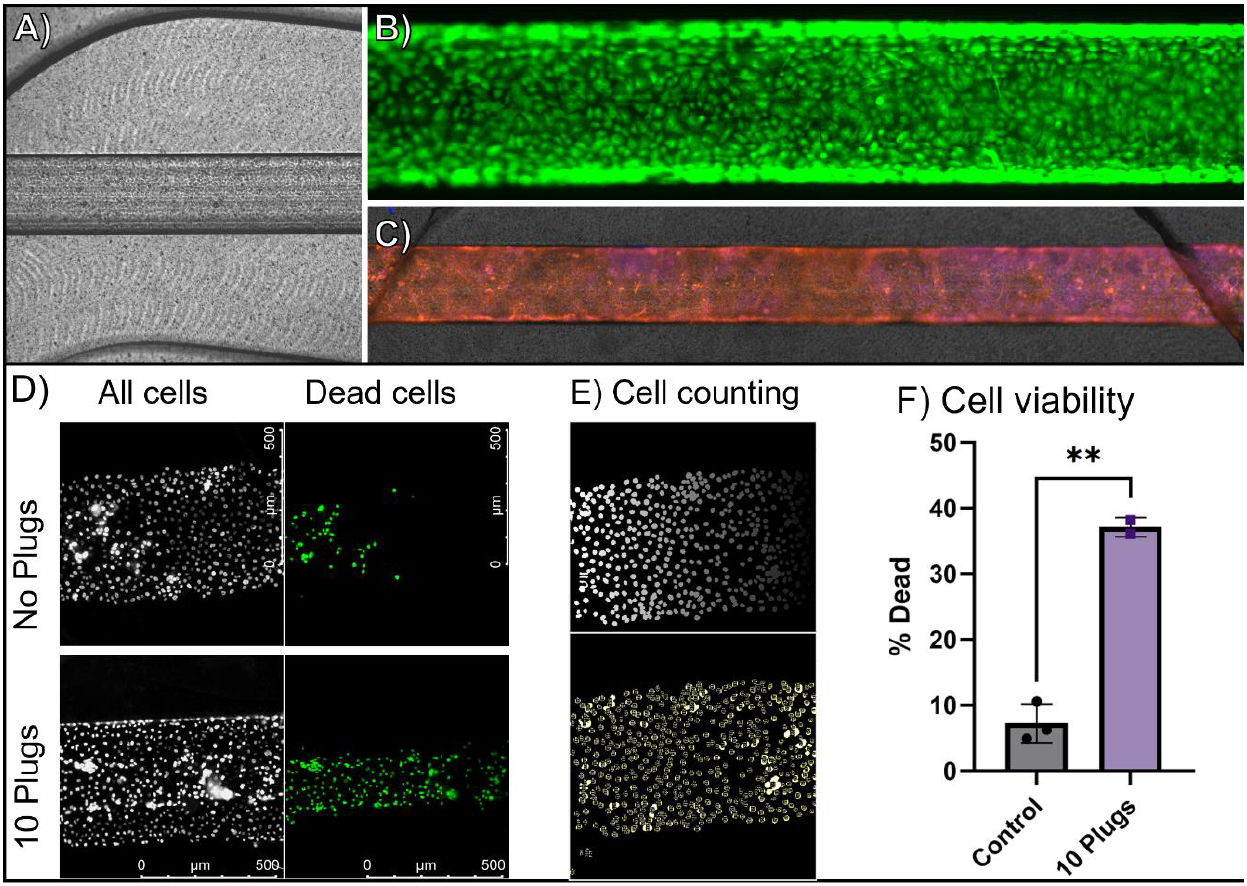
Liquid plug propagation and viscosity effects. **A)** Increasing liquid viscosity results in slower plug propagation **B-D)** Simulated plug propagation in a circular channel shows that increasing viscosity results in slower propagation speed **(B)**, matching experimental results; **(C)** comparable pressure drop across the channel but delayed rupture, and **(D)** increased maximum shear stress.

Taken together, we demonstrated a physiologically-relevant capability of our system to model viscosity effects in a microfluidic system robustly. Propagation speed relates physiologically to how much time is required to clear the fluid from an inspiratory airway. A lower propagation speed indicates that a greater portion of inspiration might be required to clear a plug. On a large scale, slower plug propagation could be expected to negatively impact overall respiratory mechanics^11^. Interestingly, amongst COPD patients, increased mucous plugs are associated with compromised lung function, more frequent exacerbations, and more severe disease^55^. Future studies in our device could investigate whether such alterations in viscoelastic liquid plug properties directly contribute to cellular pathophysiology in muco-obstructive lung diseases.

### Liquid plugs mediate cellular injury

In the final application of our model system, we demonstrated the capability to generate liquid plugs in the context of cellular injury. Future studies will take advantage of this capability to evaluate the effect of surfactant and viscosity variations on cytotoxicity. Here, we establish that surfactant-free liquid plugs are rapidly cytotoxic to exposed cells. Primary small airway epithelial cells were cultured in the microfluidic devices with a semi-circular cross-section for 3 days to reach confluence (Figure 5A). Cells remained viable as indicated by staining for live (green, calcein AM) and dead (red, ethidium bromide) cells in the channel at the end of the culture period (Figure 5B). Cells also demonstrated complete coverage of the microfluidic channel as indicated by cell membrane stain in Figure 5C (CellMask, orange; nuclei, blue). Then, either no plugs or 10 plugs were applied to the cells over a period of 50 seconds with a liquid flow rate of 100 μL/min and air flow rate of 56 μL/min (one plug every five seconds). We quantified cytotoxicity in the region where plugs were propagated, but not yet ruptured.

**Figure 5.**
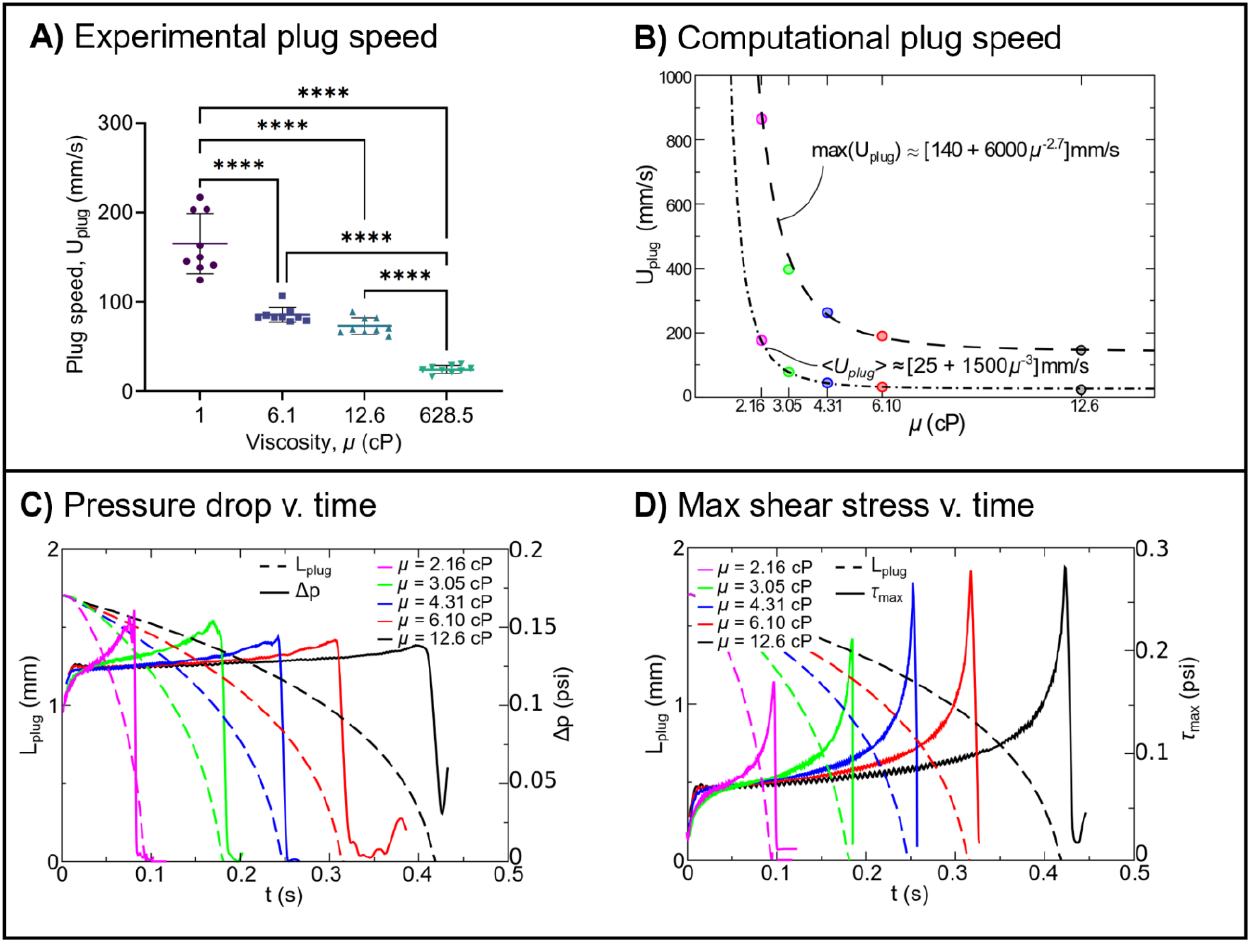
Liquid plugs are lethal to airway cells. **A)** Brightfield image of primary airway epithelial cells cultured in the microfluidic device. **B)** Live cell stain (calcein AM) shows that cells are confluent and alive 24 hours after seeding. **C)** Live cell staining for the cell membrane (CellMask) and nuclei (NucBlue) shows that cells are confluent after 3 days of submerged culture. **D)** Viability for control vs. plug condition. **E)** Example counting in CellPose (ImageJ). **F)** Plug viability for control (n=3 devices) vs. 10 plugs (n=2 devices).

Interestingly, application of surfactant-free liquid plug propagation induced epithelial cell death along the center of the channel but avoided the edges. We speculate that this cell death pattern is due to the corner sections having a thicker fluid layer than the middle due to the imperfect circularity of the channel^56^. This would result in reduced shear stresses to the corners compared to the center of the channel and therefore reduced cell death. While human airways are approximately round, irregularities in their circular geometry are observed in experimental studies. The impact of geometric irregularities, such as imperfect circularity as demonstrated here, are an interesting topic of future study.

## Conclusions

In this paper, we developed a microfluidic device and peripheral controls to model a broader range of fluid mechanical stress in the distal airways within a format that can also incorporate primary human small airway epithelial cells. We provide a robust method for leak-free custom sandwich device fabrication with a 3-dimensional semi-circular channel cross-section achieved by micro mill fabrication rather than lithography. We demonstrate the device’s ability to recapitulate both surfactant and viscosity-mediated fluid dynamics as well as the ability to capture fluid shear stress mediated cell injury. These advances significantly improve the device system capabilities for future *in vitro* studies of complex respiratory fluid mechanics and resultant cellular-level physiology.

The technical advances in microfluidic fabrication along with peripheral controllers enabled the resultant system to produce robust, highly-controlled liquid plugs including plugs with surfactant that are difficult to control due to plug instability that accompany lower surface tension, and plugs of high viscosity liquids that are difficult to control due to the high resistances to flow. We employ micro mill fabrication to achieve 3-dimensional channel design that more readily models the airway with a semi-circular cross-section rather than a high aspect ratio rectangle, as we did previously. Because micro-milled molds as well as sandwiching of porous membranes make device bonding difficult, a chemical bonding method was incorporated to enhance robustness. The semicircular cross-section enabled an interesting observation of heterogeneous liquid plug-mediated injury that depended on the cells’ distance from the centerline of the channel. This result suggests that future studies should investigate the role of airway epithelial plication (i.e., ridging) on the cytotoxicity of fluid shear stress. Additionally, the microfluidic fabrication takes advantage of a PDMS double-casting method. Micro milling the positive rather than negative mold causes increased surface roughness compared to the naïve acrylic surface, compromising the adhesive strength of the resultant PDMS components. Here, we eliminate this hurdle by milling the device as a negative onto a naïve acrylic surface. Then, we cast the positive in PDMS, creating a master mold that was then used to cast the final devices. This strategy improved the throughput of fabrication and improved the adhesion of top and bottom device layers by reducing surface roughness, although we still required chemical bonding for robustness. Finally, our cell culture method eliminates the need for pumps, tubing, and fittings during cell culture, increasing the throughput of devices and lowering costs. These improvements to the method reduce the necessary labor and costs while increasing the success rate of devices at the cell culture stage.

Future studies may take advantage of the enhanced capabilities introduced by our model system. Namely, we enable long-term studies of various types of plug exposure in a cell-culture system with primary human airway epithelium. While we have previously shown that surfactant reduces plug-mediated cytotoxicity, here we demonstrate that channel geometry also plays a role in cell injury. Importantly, human airways are heterogeneously rounded and these geometric ‘imperfections’ in circularity may impact the amount of shear stress experienced by cells. We uniquely demonstrate that imperfections in circular geometry could play a role in cellular injury, a topic that should be expounded upon in future studies. We also related the peak plug pressure to capillary pinning created under condition of high surface tension. It is important to note that static capillary resistance depends on surface wettability with more wettable (i.e., hydrophilic) surfaces generally having lower contact angle hysteresis ^57^. It is difficult to quantify the contact angles of liquids in our channel due to its curved geometry. Although PDMS is known to be mildly hydrophobic without surface treatments, our channel was exposed to multiple hydrophilic treatments. During fabrication and cell culture, the PDMS channel is exposed to oxygen plasma, UV, and protein adsorption from cell culture medium. These treatments are known to induce PDMS hydrophilicity to varying degrees^58^. We previously observed that convex shape of plugs is only achievable with hydrophilic surface coating^21^, and we similarly observe convex plugs here, suggesting that our channel is hydrophilic. High-speed movies, however, reveal noticeable contact angle hysteresis at the advancing menisci. In the human distal airways, static capillary resistance can become significant enough to prevent plug propagation, resulting in static mucus plugs. Typically this occurs under high mucous conditions, low surfactant, and/or dysregulated ion balance, such as during CF and COPD. Therefore, our model could be used to investigate the influence of mucus wettability, viscosity, and adhesiveness on the force (i.e., pressure) required to move a stagnant plug. This capability may also enable our model to serve as a platform for the evaluation of next-generation synthetic surfactants and their potential effects on static as well as dynamic capillary resistance in physiologically relevant conditions.

In summary, we present a robust, reproducible model for liquid plug generation in a leak-free porous membrane sandwiching compartmentalized microchannel device and demonstrate its potential application to several areas of pulmonary air-liquid fluid mechanical physiology. This device represents a significant advance in capabilities to study air-liquid fluid phenomena *in vitro* with the ability to manipulate liquids with a broader range of surface tension and viscosities. The platform enables future studies that will improve understanding of fluid-mediated small airway injury and ultimately lead to improved strategies for the management and prevention of fluid mechanical stress injuries to the airway epithelium.

## Supporting information

Supplemental Calculations

Device Solidworks File

## Author Contributions

Conceptualization: HV, JG, ST; Formal analysis: HV, CL, FR; Funding acquisition: ST, JG; Investigation: HV, VV, KW, CS, AL; Methodology: HV, JHL, NM; Project administration: HV; Supervision: JG, ST; Writing- original draft: HV; Writing- review & editing: all authors.

## Conflicts of interest

There are no conflicts to declare.

## Acknowledgements

This work was supported by NIH R01HL136141. HV was supported by 5T32EB006343-08. This work was also performed in part at the Georgia Tech Institute for Electronics and Nanotechnology, a member of the National Nanotechnology Coordinated Infrastructure (NNCI), which is supported by the National Science Foundation (Grant ECCS-2025462). We thank Dr. Aaron Lifland and Georgia Tech’s Optical Microscopy Core for lending their Olympus microscope for this work. Additionally, we acknowledge Dr. Octavian Ioachimescieu, Dr. Rabindra Tirouvanziam, and Dr. Jocelyn Grunwell for their insight and expertise supporting this project. Figures were created with Biorender.

## Notes

### Competing Interest Statement

The authors have declared no competing interest.

