## Supplemental Figures for "Liquid plug propagation in computer-controlled microfluidic airway-on-a-chip with semi-circular microchannels"

### Slide 1
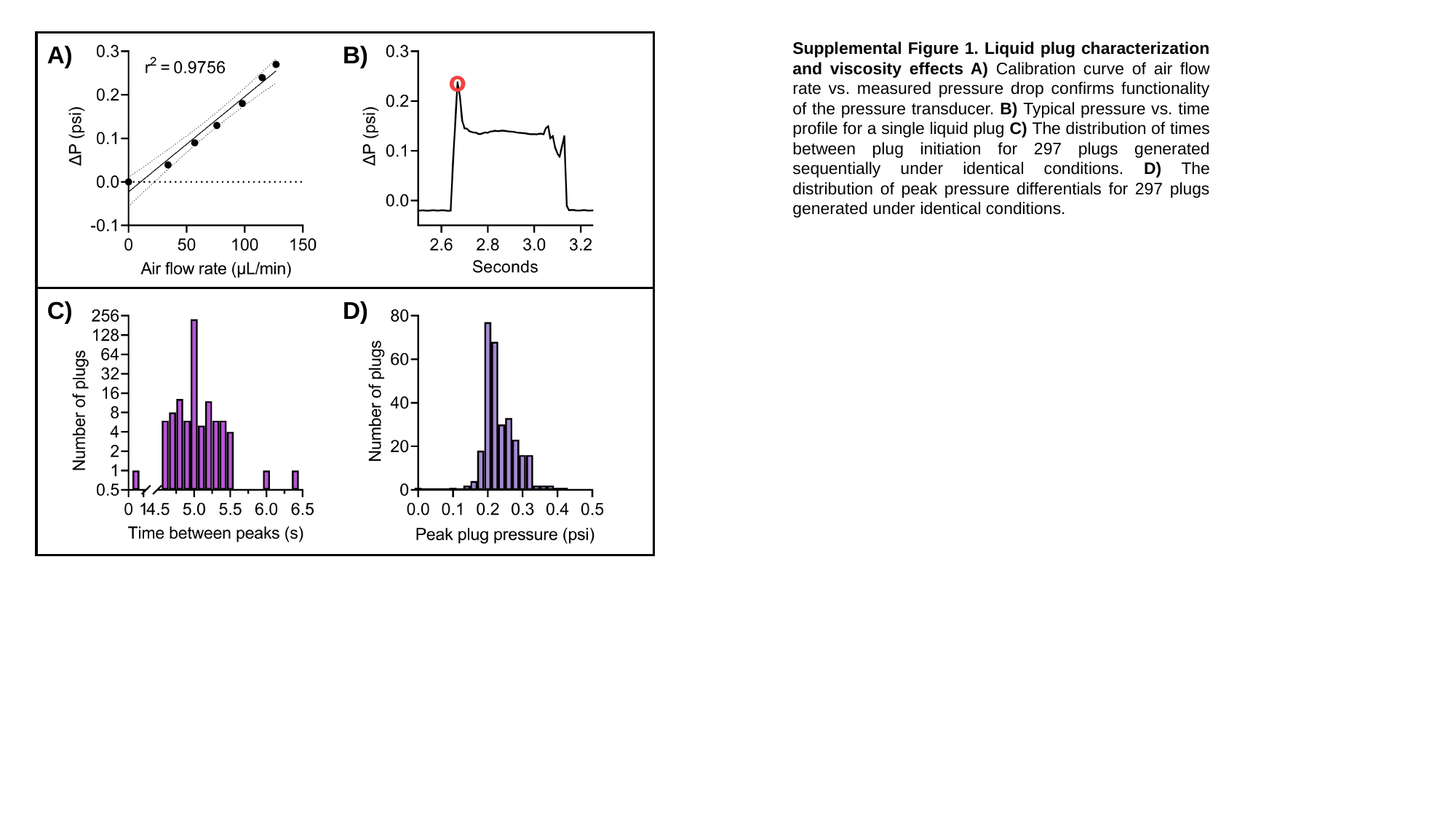

Supplemental Figure 1. Liquid plug characterization and viscosity effects A) Calibration curve of air flow rate vs. measured pressure drop confirms functionality of the pressure transducer. B) Typical pressure vs. time profile for a single liquid plug C) The distribution of times between plug initiation for 297 plugs generated sequentially under identical conditions. D) The distribution of peak pressure differentials for 297 plugs generated under identical conditions.
A)
B)
C)
D)

### Slide 2
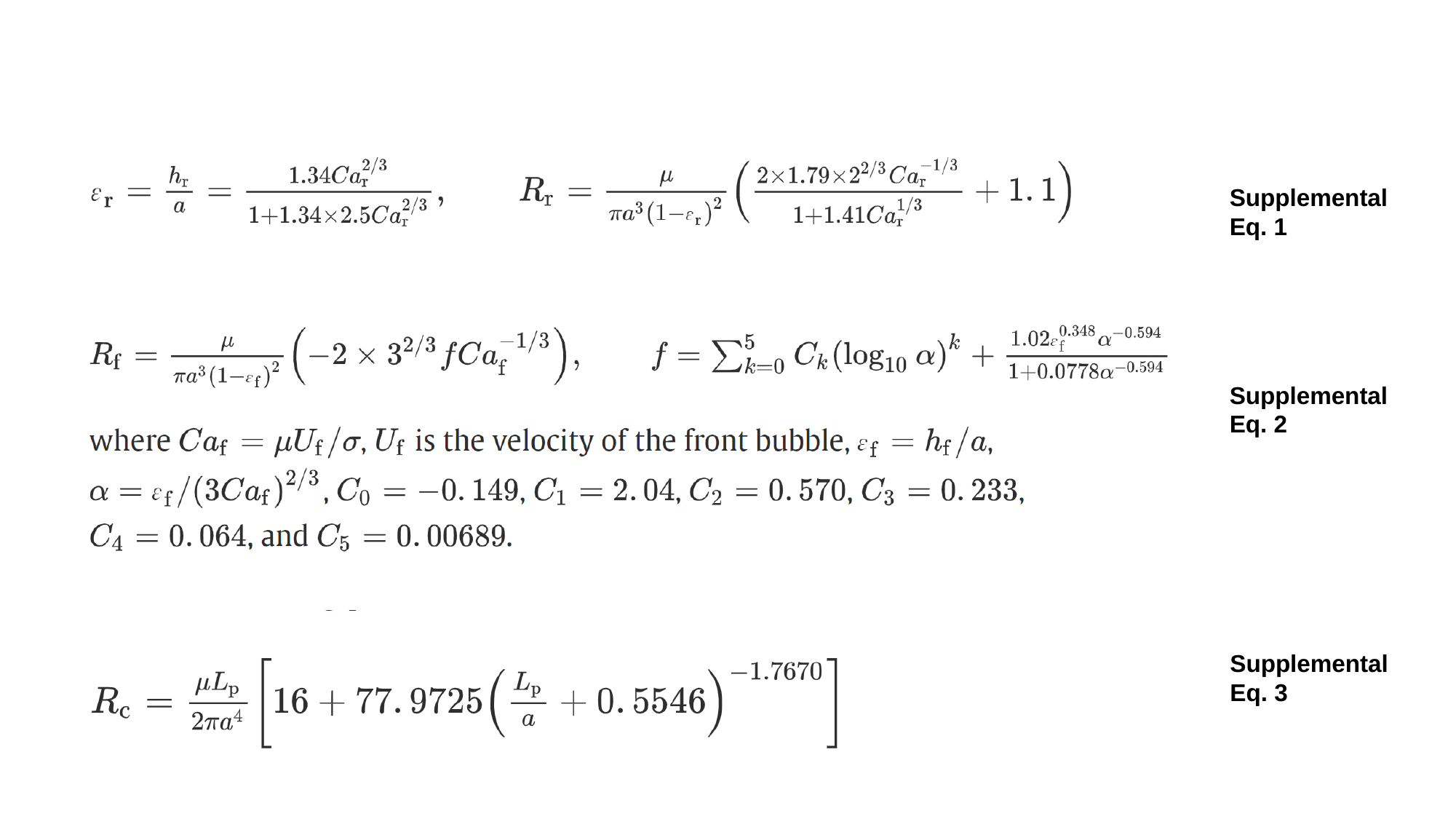

Supplemental Eq. 1
Supplemental Eq. 2
Supplemental Eq. 3
